# Loading effects on the performance of needle free jet injections in different skin models

**DOI:** 10.1101/853481

**Authors:** Pankaj Rohilla, Idera Lawal, Andrew Le Blanc, Veronica O’Brien, Cormak Weeks, Whitney Tran, Yatish Rane, Emil Khusnatdinov, Jeremy Marston

## Abstract

Intradermal delivery of vaccines with jet injection is one of the leading alternatives to conventional delivery with hypodermic needles via the Mantoux technique. However, for a given fluid, the effects of various parameters related to injector design, as well as skin properties are still not well understood. Whilst the key design parameters are orifice diameter, jet speed, ampoule volume, and standoff distances, we must also consider applied load of the device on the skin, and axial skin tension. These parameters are all studied herein using different ex-vivo models (guinea pig, pig and human skin) and different fluid viscosities. We find that the applied load can have a significant effect on the amount of drug delivered through the skin, as well as the fluid dispersion pattern in the intradermal tissues. Regardless of skin type or fluid viscosity, we show that minimal standoff and applied loads of approximately 1 kg should be used to maximize injection efficiency when targeting intradermal tissue.

## INTRODUCTION

Needle-free injector devices use impulse force for targeted vaccine delivery into the intradermal, subcutaneous and intramuscular regions. Different actuation mechanisms such as spring force [1], Lorentz coil [2], compressed gas [3], piezoelectric motors [4] and laser induced cavitation [5, 6] have been used in the development of the needle free jet injectors, where the working principal is creation of a high-speed microscale jet that can puncture the skin and deposit drug into tissues beneath. To date, the clinical use of such devices has been limited possibly due to a confluence of cost, pain, bruising and inefficient delivery of the vaccine. However, the advent of high-immunogenicity DNA-based vaccines and the need for high force to deliver such viscous products has increased interest in improving the design of jet injector devices. It is also noted that jet injections result in higher absorption rates as compared to needle injections due to the formation of more diffused dispersion patterns inside the skin [1].

The major challenges in development of the jet injectors are inefficient delivery (percentage of expelled drug actually delivered across the skin) and controllability of the depth, which is especially relevant in targeted vaccine delivery. To combat this, various design parameters such as orifice diameter, ampoule volume, standoff distance, viscous losses in the nozzle, and jet speed profile have been studied in the literature [2, 7, 8]. The physical properties of the vaccine and mechanical properties of the skin also affect the delivery efficiency and the dispersion of the drug inside the skin [9, 10].

In order to inject vaccine via a jet injector, an individual has to keep the jet injector placed perpendicular on the skin. A spacer ring has previously been used [11] to achieve this, which has the dual effect of implementing a stand-off distance between the orifice and skin. However, in the act of keeping the jet injector on the skin, a force of different magnitude is applied, depending on the individual administering the injection. The nature of the applied force is both compressive and tangential, which results in additional tension across the skin and stress in the tissue beneath the skin.

Whilst the effect of this applied load on transdermal drug delivery has not been systematically investigated before, we note that needle-free injection devices have been designed specifically to pre-tension the skin using either a threshold applied normal load [12], or a nozzle device to stretch the skin axially before actuating the jet [13]. In this article, we have therefore attempted to quantify the effect of applied loading on the performance of the jet injections in addition to other parameters such as standoff distance, liquid viscosity, ampoule volume, orifice diameter, skin type (animal) and skin support (underlying tissue).

The availability and cost of human skin limits its use in drug delivery studies. Thus, various skin models such as rodent and mammalian skin have been used to mimic human skin behavior in the literature [14]. Porcine skin is the closest skin model of human skin after a freeze-thaw cycle [15]. In this study, we have used porcine skin, guinea pig skin and human skin to inject liquids with jet injectors. Delivery efficiency and aspect ratio of dispersion pattern formed by the liquid under the skin were used to characterize performance. Injections into skin on porcine cadavers as well as injections into freshly excised and frozen skin parts of the same cadavers were also performed in this study.

We have both qualified and quantified the effect of a wide range of variables associated with the skin, injector device and the fluid viscosity on the jet injection delivery. The results presented aid in better understanding of these parameters to overcome the current challenges in the development of needle free jet injector devices.

## METHODS

### Skin samples

Porcine, Guinea pig skin and human skin were procured from animal sciences (Texas Tech University), Inovio Pharmaceuticals and National Disease Research Interchange (NDRI), respectively, and all had thickesses around 3-5 mm. Guinea Pig skins were kept in a freezer at −4 °C whereas human skin and porcine skin were stored in a freezer at −20 °C. In addition to injections performed on cadavers, the porcine skin was harvested from Yorkshire-Cross pigs euthanized at 13 weeks of age. Human skins used were from Caucasian males and females in the age range of 55-81 and with BMI indexes ranging from 21.50 to 29.86 (see table S14).

### Skin supports

In order to mimic different tissues beneath the skin samples, we used either (a) rigid glass substrate, (b) 1 cm lean porcine tissue, (c) 2 cm lean porcine tissue, or (d) 1 cm porcine fat layer. All porcine tissue used for supporting human and GP skin was procured from a local butcher and was frozen at −4 °C. Skin and pork meat was thawed to room temperature before the injection.

### Device and loading protocol

A spring-powered device (Bioject ID Pen) was used to perform jet injections, which ejects a volume, *V*, of either 50 *µ*l or 100 *µ*l. The orifice diameter, *d*_*o*_, was either 155 or 175 *µ*m and the stand-off, *S* was either 0, 2 or 14 mm. DI water and 80% glycerol were used as injectate liquids due to their viscosity gap of nearly 2 orders of magnitude with *µ* of 1 mPa.s and 84 mPa.s, respectively. Once loaded with a filled cartridge, the jet injector was firmly fixed onto a vertical stage with uniaxial motion, and lowered onto the skin sample, which was kept on a mass balance, to register the applied normal load. The value of normal load could then be varied simply by adjusting the vertical position of the injector. In addition, a miniature load cell (Futek - LLB 130, 50 lb, Item #FSH03880) was embedded within the mass balance to provide more detailed measurement of the injection force at a sample rate of 4800 sps (see figure 1(a)). Applied normal load, *L*_*n*_, on the skin was observed visually from the mass balance, and was varied from 0 *–* 2 kg. To achieve axial loading, *L*_*a*_, skins were sewed on two sides with one side fixed on a post and the other side attached to a hanging load of either 0.5 kg or 1 kg. To avoid structural damage to the skins, axial loading was limited to the maximum hanging load of 1 kg.

**Figure 1:**
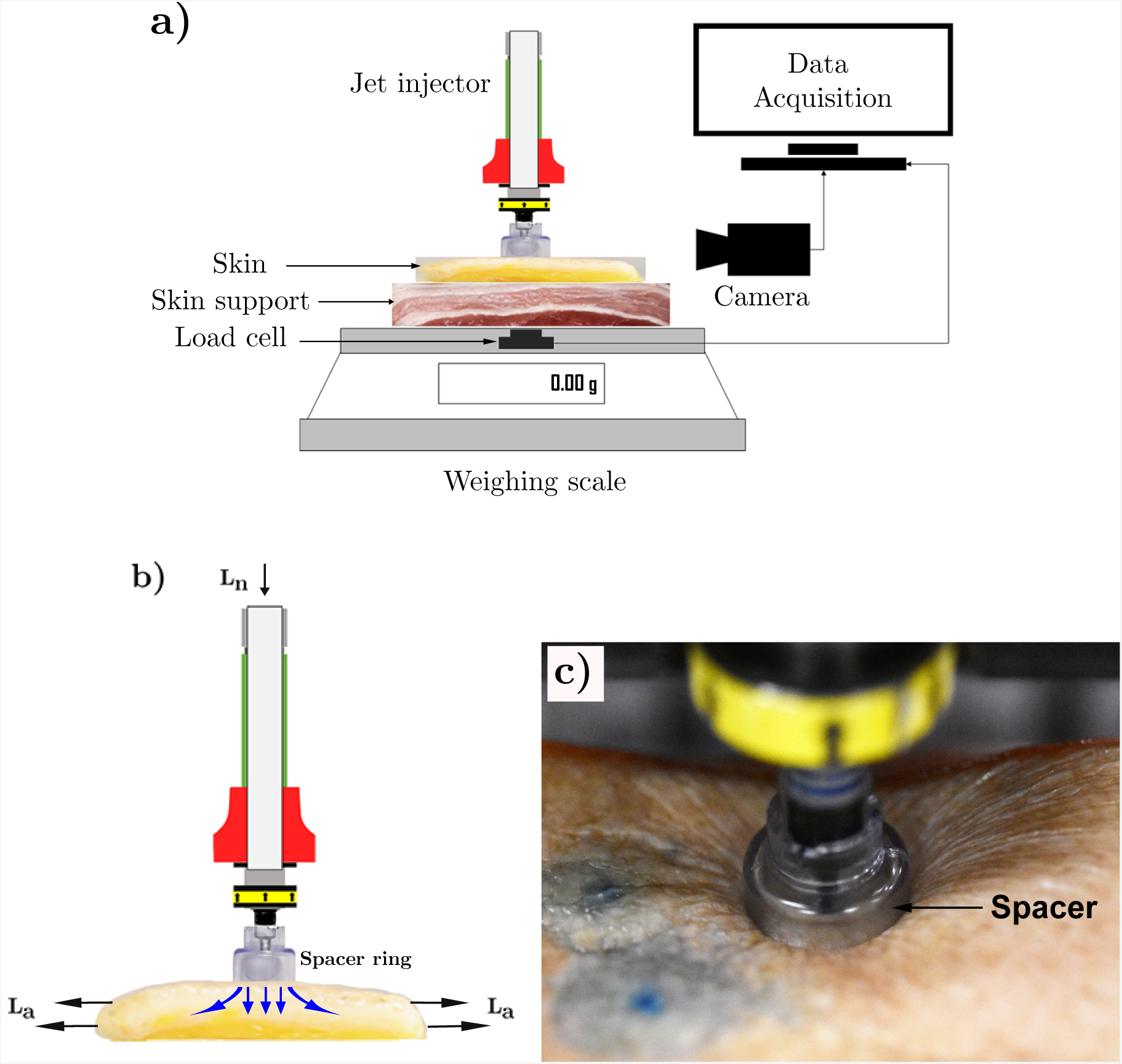
Experimental. **a).** Schematic of the experimental setup **b).** Schematics of applied normal load, *L*_*n*_ and axial load, *L*_*a*_, both shown by black arrows, and hypothesized distribution of stress in the skin tissue shown by blue arrows and **c).** Human skin under tension by a spacer ring with an applied load of 1 kg by jet injector.

### Imaging and characterization

To visualize the dispersion of liquids into the skin, Trypan Blue (Sigma Aldrich) was added to the liquids used for the injection in concentration of 1 mg/ml. After injection, skins were frozen to −4 °C, and then cut along the cross-section of the injection site to visualize the intradermal bleb, whose dimensions (total depth, *h*, width, *w*, and depth at maximum width, *d*) were measured using image processing in Matlab (see figure 3 for an example). We then characterize the dispersion pattern with aspect ratio *AR* = *d/w*. To characterize efficiency of the delivery, *η*, we used a volume-based measurement, given by the ratio of fluid injected across the skin to that ejected by the device as:

**Figure 3:**
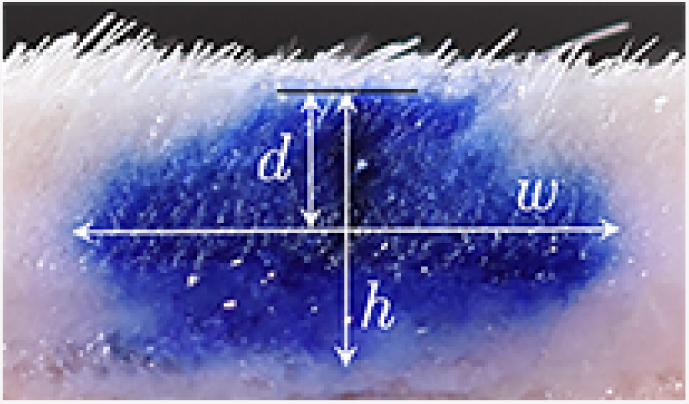
Dimensions of the bleb formed inside the skin after jet injection: w is the maximum width, h is the total depth of the bleb from the surface of the skin, and d is from the skin surface to the depth at maximum width.

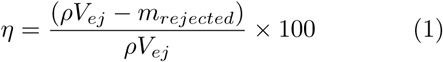

For each individual configuration of human skin, a minimum of 10 replicates were done, whereas for guinea pig and porcine skin, a minimum of 5 replicates were done. Statistical significance of the different parameters (e.g. *L*_*n*_, *L*_*a*_, *µ, d*_*o*_, *V, S*, skin type etc.) on both *AR* and *η* was determined by ANOVA tests with significance level *α* = 0.01.

## RESULTS AND DISCUSSION

### i. APPLIED LOADING

Skin is a viscoelastic material which shows relaxation when stretched and kept at a constant strain [16]. Pressing a jet injector with a spacer ring results in a compression force on the skin as well as tangential stress within the skin, as shown in figure 1(b). The behavior of the skin has been seen to be dependent on the load history; Cyclic loading in a non linear fashion yields a shift in stress-strain curves until a convergence [16, 17]. We therefore also studied the effect of load history by using three different loading mechanisms.

In loading mechanism type *A*, the jet injector was initially pressed on the skin with an extra ≈40-50% of the target load (i.e.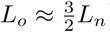). As observed from the weighing balance, the load on the skin decreases non-linearly due to the skin relaxation in which applied load distributes itself within the skin layers. An example of the loading profile, as determined by the high-resolution load cell, for type *A* mechanism for *L*_*n*_ ≈ 500 g on guinea pig (GP) skin is presented in figure 2. Here, the injector was loaded onto the skin at a normal load of ≈710 g corresponding to a force of ≈7 N. Applied load decreases due to the skin relaxation and the jet injection was actuated once the load reaches a value of ≈500 g. Injection duration can be seen in the figure as a spike in the force. Although the increase in force during jet injection was observed to be the order of ≈4 N, this is due to the extra manual force exerted during triggering; The actual force profiles for jet injections (without any manual force) with similar nozzle and liquid parameters was reported in our past work [18] and is of order of 1 N.

**Figure 2:**
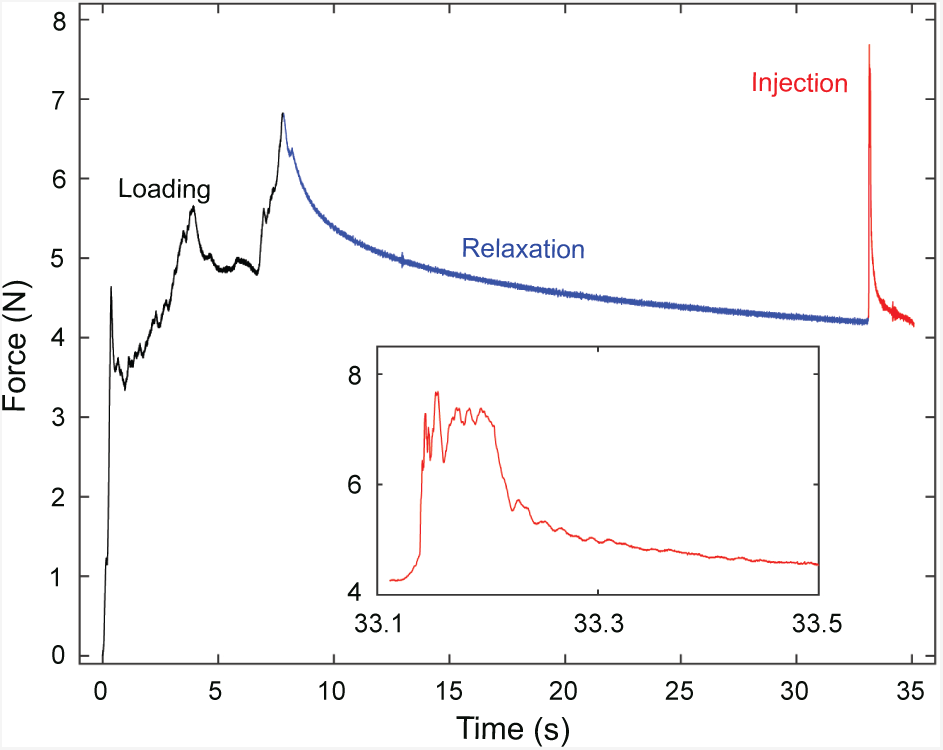
Force measured by the load cell underneath the weighing balance for 80% glycerol injection through a nozzle with orifice diameter of 155 *µ*m into GP skin with S = 0 mm and *L*_*n*_ = 0.5 kg for the entire duration including (1) loading stage, (2) relaxation stage and (3) jet injection stage. Inset shows force profile for jet injection duration.

In type *B* mechanism, skin was loaded by extra ≈50% weight followed by relaxation to the required weight for 2 cycles. The injection was then triggered when the weight approaches the required loading value in the second cycle. Lastly, Type *C* mechanism is identical to type *A* mechanism with the exception of initial loading weight (*L*_*o*_), which was chosen to be twice as that of the required value of the load i.e. *L*_*o*_ = 2*L*_*n*_.

### ii. LIQUID DISPERSION

The depth and shape of the bleb (i.e. fluid distributed within the intradermal tissue) ultimately dictates the effectiveness of delivery and the diffusion of the vaccine within the skin. Liquid dispersion inside the skin was characterized by the aspect ratio of the bleb formed in the skin, i.e. *AR* = *d/w*, as shown graphically in the example in figure 3. One would intuitively expect fluid dispersion in the intradermal tissue to be affected by various parameters such as viscosity and mechanical properties of the skin (e.g. skin type and applied load). However orifice diameter and stand-off distance affect both the impact velocity and impact footprint of the liquid jet [18], and therefore it is instructive to examine these effects as well. As a brief overview, cross-sections of blebs formed after jet injection into the different skin types, under similar conditions are presented in figure 4.

**Figure 4:**
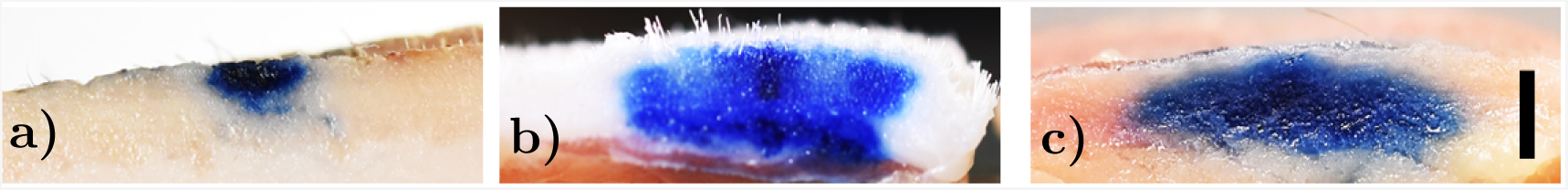
Cross sections of blebs for water injected into (a) Porcine skin, (b) guinea pig skin and (c) human skin. All injections correspond to *d*_*o*_ = 155 *µ*m, *S* = 0 mm, and *L*_*n*_ = 1 kg. *Scale bar represents 5 mm.*

The effect of increasing applied load caused by the hollow cylindrical spacer creates an extruded dome shape on top of the skin, which becomes more defined with increased load, as can be seen in figure 5. However, the bleb shape under the skin also changes from an oblate ellipsoidal to cylindrical shape with increase in the normal static load, as seen in figure 6. That is, the bleb width increased as the applied normal load was increased to 2 kg. This can be understood by considering the compressed state of the skin when under load; the compressive stress in the skin and intradermal tissues leads to axial bias for dispersion, resulting in fluid being squeezed farther in the horizontal direction.

**Figure 5:**
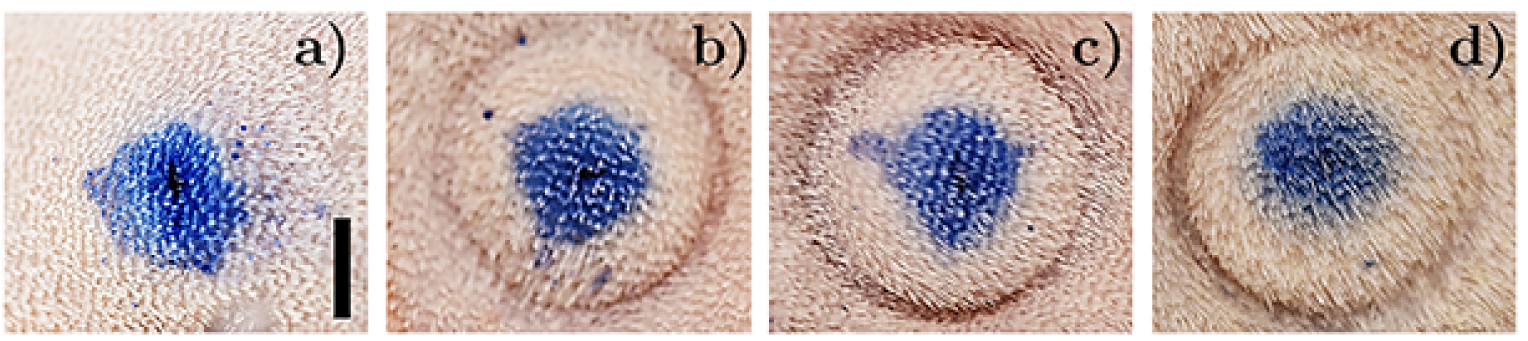
Effect of applied load on the top of the GP skin supported by 2 cm thick lean pork after jet injection of 100 *µ*l dyed water through a nozzle with orifice diameter of 155 *µ*m for *L*_*n*_ of: a. 0 kg, b. 0.5 kg, c. 1 kg and d. 2 kg. *Scale bar represent 5 mm.*

**Figure 6:**
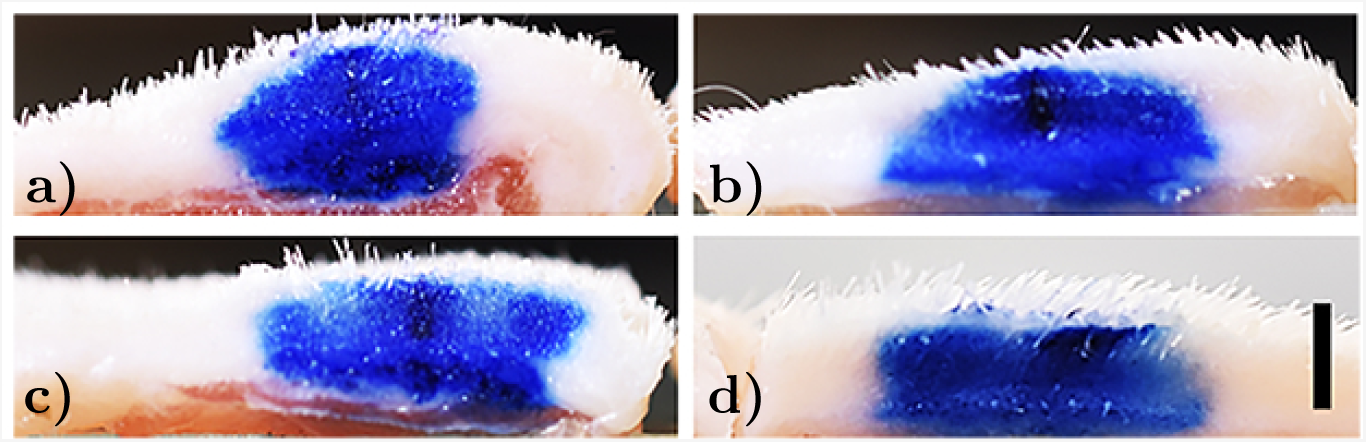
Cross sections of the bleb formed in GP skin supported by 2 cm thick lean pork injected by 100 *µ*l dyed water through a nozzle with orifice diameter of 155 *µ*m for *L*_*n*_ of: a. 0 kg, b. 0.5 kg, c. 1 kg and d. 2 kg. *Scale bar represents 5 mm.*

In addition to the static load, the effects of standoff and fluid viscosity assessed previously [10] are also hypothesized to become important due to their respective influences on the jet impact footprint and diffusion coefficient in porous substrates. Indeed, increasing standoff distance from 0 mm to 14 mm also showed transition to cylindrical shape of the skin bleb. However, blebs formed from 14 mm standoff were narrow in width with higher aspect ratio as compared to injections from zero standoff distance. A comparison of aspect ratio, *AR*, of the blebs formed inside the GP skin for different standoff, applied normal load and injectate viscosity is presented in figure 7, where the most significant variation occurs for *L*_*n*_ = 1 kg, but in all cases, increasing the standoff distance increased *AR* accordingly.

**Figure 7:**
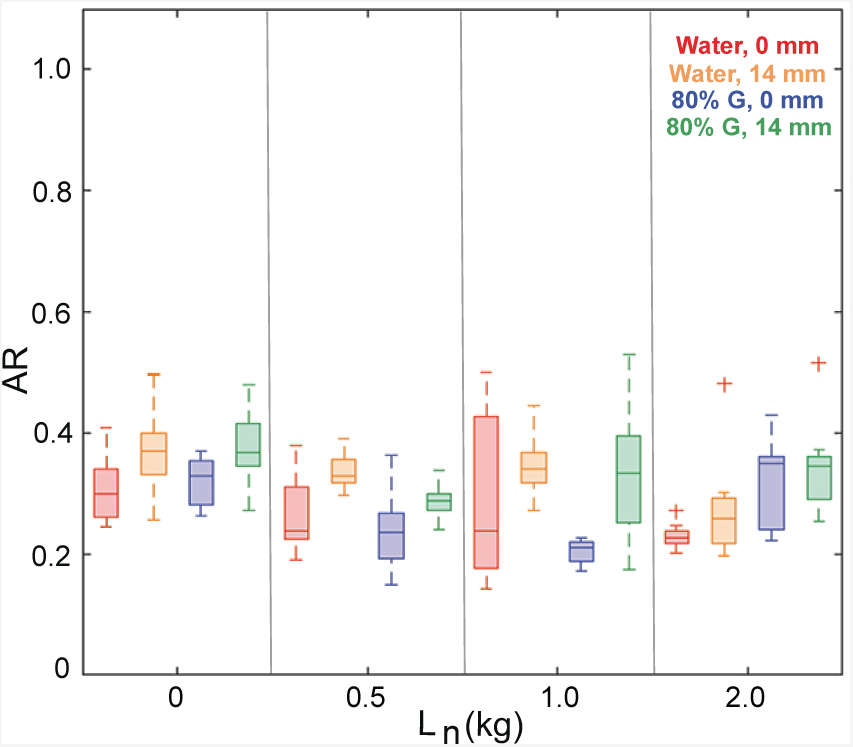
Comparison of aspect ratio of the blebs formed in the GP skin supported by 2 cm thick leaned porcine meat for different *L*_*n*_, *s* and *µ* and *d*_*o*_ = 155 *µ*m.

Complete analysis for all skin types shown in supplemental figures S3, S4, and S5 corresponding to guinea pig, porcine, and human skins. However, two key observations that must be highlighted are for the porcine skin (figure S4), where we were able to compare freshly excised skin to skin having undergone a single freeze-thaw cycle: For injection into ventral region skin, the bleb shape changed significantly from fresh (*AR* ≈ 0.5) to frozen-thawed (*AR* ≈ 0.24). However, the same injections into dorsal skin were invariant to the freeze-thaw cycle with *AR* ≈ 0.1 in all cases (see figure S4b). Furthermore, for the freshly excised skin, the effect of orifice diameter was also significant for both viscosities (see figure S4a), where *AR ≈* 0.25 for *d*_*o*_ = 155 *µ*m, but *AR ≈* 0.1 for *d*_*o*_ = 175 *µ*m.

### iii. DELIVERY EFFICIENCY

#### A. PORCINE SKIN

In this study, we have used three different states of the porcine skin including cadaver, freshly excised and frozen skin. Liquid was injected into the cadavers within an hour of euthanasia such that injections were performed with minimum alteration in skin mechanical properties. Furthermore, skin from different anatomical parts including ventral and dorsal parts was also excised and used both before and after a cycle of freezing and thawing. Mechanical properties of porcine skin after a single cycle of freezing and thawing are nearly same as that of human skin [15]. However ventral porcine skin is known to be more anisotropic in comparison to the dorsal part [19]. Although various properties of porcine and human skin are similar such as morphology, immunogenicity, physiology and composition of cells; mechanical properties of the top layer of the skin, i.e. stratum corneum (SC) is different. Porcine SC exhibits higher Young’s modulus (*E*_*SC*_) than human skin, which increases further after a freeze-thaw cycle whereas a decrease in the *E*_*SC*_ was observed for human skin after a freezing cycle [15].

##### LOAD

No significant effect of applied load was observed on the delivery efficiency of the injectate inside the freshly excised porcine skin (*p >* 0.05 as per table S9). However, large intra-sample variation (*η* in range of 11%-53%) was observed for the normal load of 1 kg as shown in figure 8.A. The delivery efficiency of the jet injection was calculated on the basis of liquid left on the top of the skin after the jet injection, as per equation (1).

##### VISCOSITY

For water injected into fresh porcine skin, the delivery efficiency was *η* ≈ 20 – 40 %, as shown in figure 8.A. However, by increasing the fluid viscosity (by nearly an order of magnitude) - we observe lower *η* for 80% glycerol in comparison to that obtained with DI water for an applied normal load of 1 kg.

##### STANDOFF DISTANCE

Three different standoff distances (*S* = 0, 2, 14 mm) between the nozzle orifice exit and the fresh dorsal skin were used. A reduction in the percentage delivery was observed with increase in the standoff distance from 0 mm to 2 mm as shown in figure 8.A. Whereas the best overall performance in terms of highest efficiency was observed for *S* = 0. There was large intra-sample variation in the data for standoff distance, rendering the effect of *S* on *η* insignificant (*p >* 0.05) for fresh skin.

**Figure 8:**
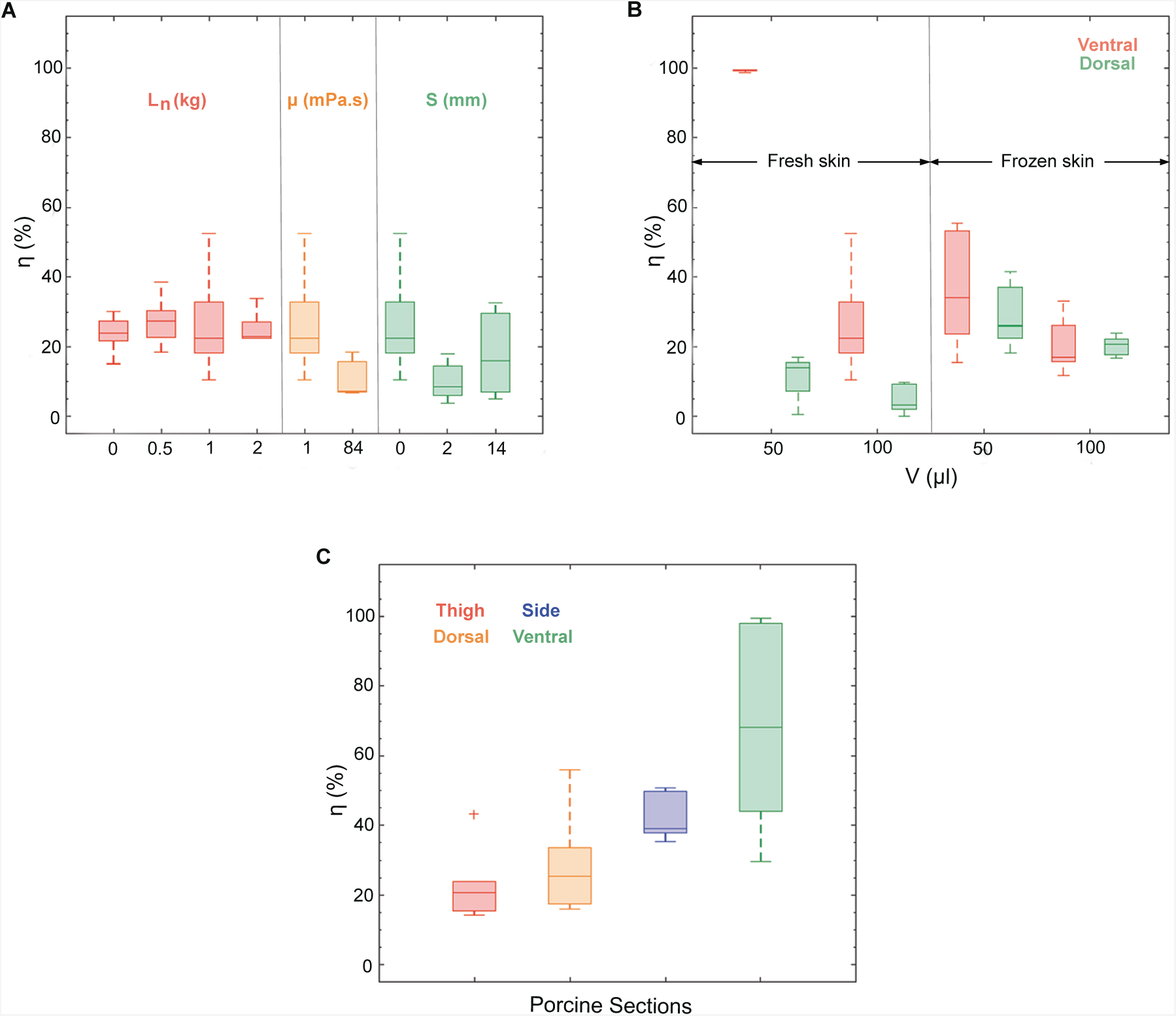
Injection efficiency (*η*) of jet injections into porcine skin. **a.** Effect of normal static load (*L*_*n*_), liquid viscosity (*µ*, mPa.s) and standoff distance (*S*, mm) on delivery efficiency of the liquid in fresh excised porcine skin (* represents *d*_*o*_ = 175 *µ*m, otherwise *d*_*o*_ = 155 *µ*m). **b.** Delivery efficiency of dyed water in porcine skin from ventral and dorsal regions affected by the state of the skin, thickness and liquid volume after a single freeze and thaw cycle for *d*_*o*_ = 155 *µ*m and *S* = 2 mm. **c.** Delivery efficiency of dyed water injection in the porcine cadaver into thigh, dorsal, side and ventral regions for *d*_*o*_ = 155 *µ*m and *S* = 2 mm

##### STATE OF THE SKIN

The anatomical location of the skin, and its state (fresh or frozen-thawed) are important for determining mechanical properties of the skin, and therefore should affect the volume which can be delivered within the skin. A summarized comparison is presented in figure 8.B, where we found nearly complete delivery (*η* ≥ 95%) was achieved for injection of *V* = 50 *µ*l water in the freshly excised skin from the ventral region. However, efficiency decreases significantly for *V* = 100 *µ*l to *η* ≈ 20 – 30%. In contrast, delivery efficiency in the dorsal skin was low in both cases (*η* ≈ 15% and 5% respectively). Porcine skin showed higher delivery efficiency in the dorsal part of the skin after a cycle of freezing and thawing. On the other hand, ventral part of the skin after a freeze-thaw cycle showed lower injection efficiency in comparison to the fresh skin. Nearly complete delivery of 50 *µ*l water in freshly excised skin from ventral part is an exception case of high percentage delivery. This exception can be attributed to the highly localized variability in the mechanical properties of the porcine skin. In contrast, the higher delivery in frozen porcine skin from dorsal region can be attributed to higher diffusivity as compared to the fresh skin. Increase in diffusivity of the skin from fresh to frozen is due to the increase in porosity from structural damage as the freezing induces formation of the ice crystals within the skin [20].

Porcine cadavers were also injected with water into different parts corresponding to the different mechanical properties (Thigh, Dorsal, Side, Ventral). Effect of variation in anatomical parts on the injection delivery efficiency was very significant (*p <* 0.05), with higher efficiency observed for the ventral skin followed by side skin, dorsal skin and thigh skin as shown in figure 8.C. Thus, percentage delivery via jet injections was observed to be inversely proportional to stiffness of the porcine skin.

#### B. GUINEA PIG SKIN

Guinea pig skin was also used as a surrogate for human skin. For these trials, we also used axial loading (*L*_*a*_) in addition to normal loads (*L*_*n*_) of 0 kg, 0.5 kg, 1 kg and 2 kg. Variation in the thickness of subcutaneous and muscular layers of the skin was mimicked by supporting the GP skin with leaned porcine meat of different thickness of 1 cm and 2 cm and a fat layer of 1 cm in thickness. Also, a rigid support (glass slab) was used to mimic the skin supported by the bone in case of axial loading experiments.

As shown in figure 9.A, GP skin supported by 2 cm porcine meat exhibited significant effect of applied normal load on the delivery efficiency of water and 80% glycerol (*p <* 0.05) for standoff distance of 0 mm. However, effect of *L*_*n*_ was insignificant (*p >* 0.05) for jet injection from the standoff distance of 14 mm for different injectates. A normal load of *L*_*n*_ = 1 kg resulted in the highest delivery efficiencies with *η* ≥ 95 % for water and *η* ≈ 75% for 80% glycerol, as shown in figure 9.A. However, further increasing the normal load to *L*_*n*_ = 2 kg resulted in lower delivery efficiency of both liquids. A similar effect of *L*_*n*_ on injection efficiency was observed for the GP skin on different underlying supports (see figure S2). Again, we can interpret this result in the context of tension and stress within the intradermal and underlying support tissues; for low loads (*L*_*n*_ *<* 0.5 kg), the underlying support is still pliable so that it can deflect due to the impulsive action of the jet. At high loads *L*_*n*_ ≥ 2 kg, the underlying support is compressed and stiff, and the intradermal tissues are also under significant compressive stress, resisting liquid dispersion throughout the injection. Therefore, the value *L*_*n*_ ≈ 0.5 − 1 kg represents a near-optimal trade-off between these two regimes which removes the influence of underlying tissue, but also correctly tensions the skin for puncture and liquid dispersion.

Axial load was applied in addition to the normal static load to understand the effect of additional tangential forces within the skin. Figure 9.B shows the effect of different magnitudes of loading on the injection efficiency of the dyed water inside the GP skin supported by glass or leaned pork. With fixed axial loading (*L*_*a*_) of 0.5 kg on GP skin supported by 2 cm thick lean pork, effect of *L*_*n*_ on *η* was significant (*p <* 0.05). However, when compared to results in figure 9.A, it is clear that addition of axial load is detrimental to injection efficiency, since 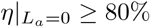 but 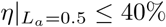. We propose that axial load in addition to normal load results in too much stress within the skin that jet pressure cannot overcome.

In case of higher axial load of 1.0 kg, GP skin is even more highly tensioned and addition of normal load resulted in lower injection efficiency. Overall effect of axial loading with constant normal static load either showed insignificant effect or a lower injection delivery. Therefore, the results for GP skin advocate that an optimal configuration exists. For GP skin, this configuration is: *S* = 0 mm, *L*_*n*_ = 1 kg, *d*_*o*_ = 155 *µ*m, and 2 cm thick underlying tissue.

**Figure 9:**
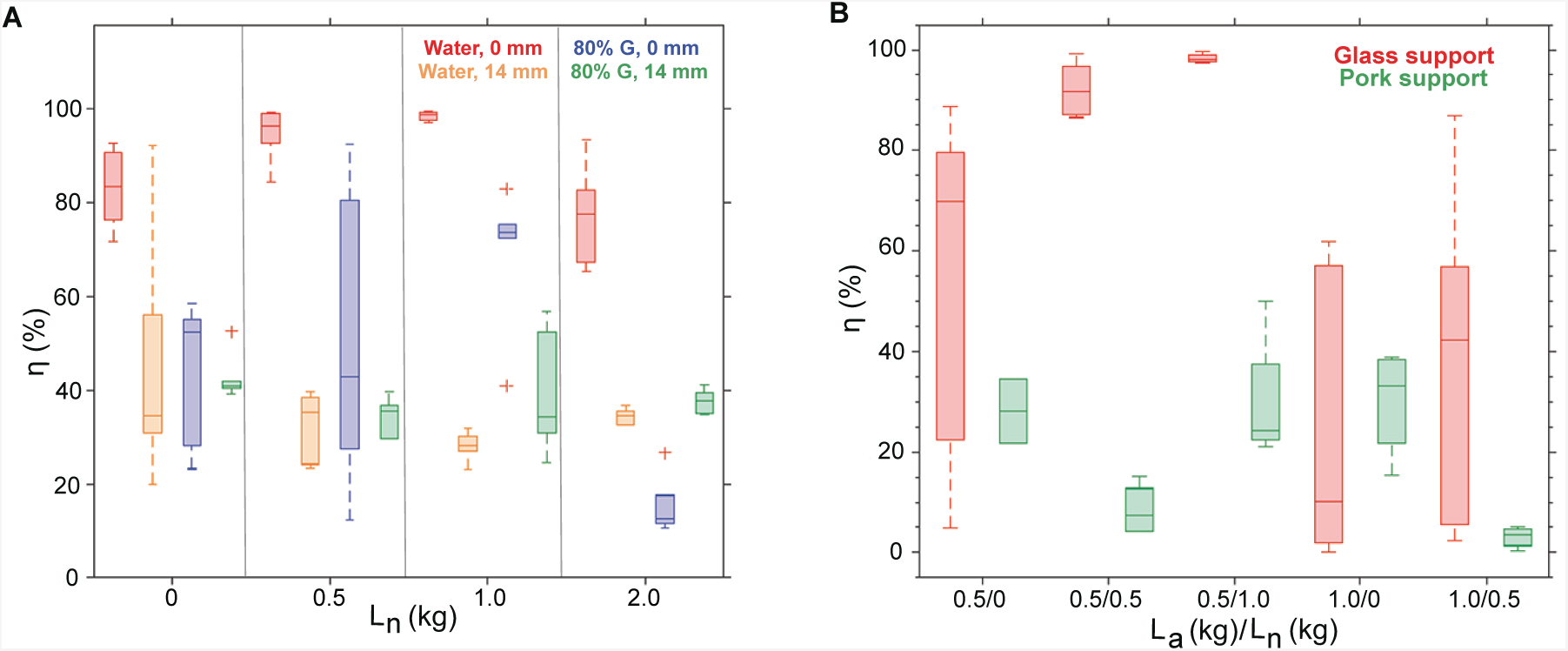
Injection efficiency (*η*) of jet injections in guinea pig skin. **a.** Effect of normal load (*L*_*n*_), standoff distance (*S*) and injectate viscosity (*µ*) on *η* for GP skin supported by 2 cm thick lean pork **b.** Effect of axial load (*L*_*a*_), normal load (*L*_*n*_) and skin support on *η* for dyed water injection, *S* = 0 mm and *d*_*o*_ = 155 *µ*m.

#### C. HUMAN SKIN

Human skin varies in terms of mechanical properties with age, loading and for different anatomical regions [21, 22, 23, 24, 25]. Elastic modulus of human skin increases with aging in addition to significant variation in thickness, stiffness and tension [26]. Behavior of human skin is visco-elastic in nature and the mechanical response to loading depends on the support or the backing material. After loading, relaxation occurs faster for skin over the muscle in comparison to the skin supported by the bone. Also, skin over the muscle can tolerate higher applied load as compared to the skin supported by the bone [27]. We have used porcine tissue and glass slab as the backings underneath human skin to understand the effect of stiffness of the skin support on jet injections with applied load. Different loading mechanisms, standoff distances and ampoule volumes were used to compare the efficiency of injections for a constant normal static load of 1 kg as shown in figure 10.A.

**Figure 10:**
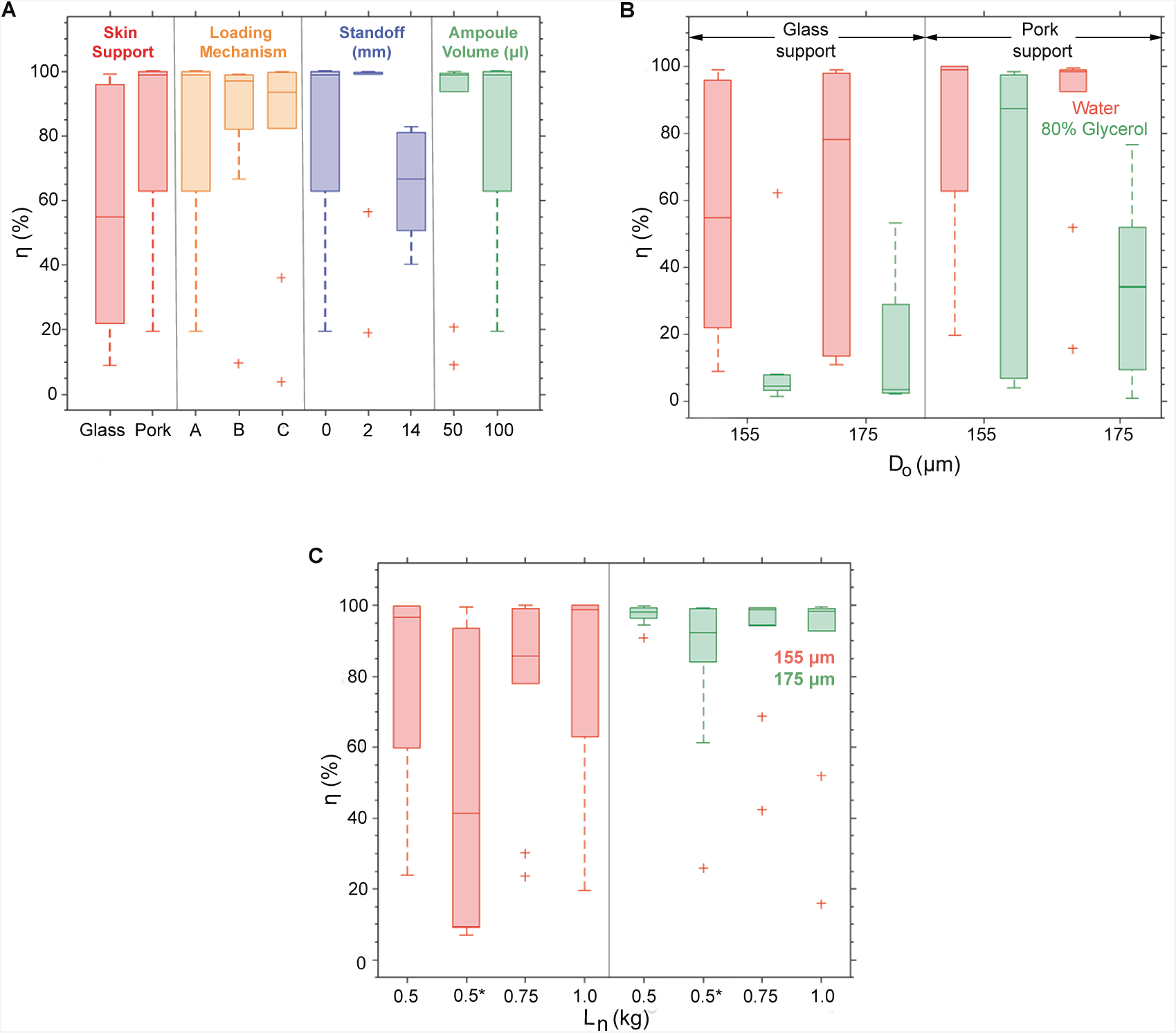
Injection efficiency (*η*) of jet injections of water into human skin. **a.** Effect of skin support, loading mechanism, standoff and *V* on *η* for *L*_*n*_ = 1 kg. **b.** Effect of *d*_*o*_, skin support and injectate viscosity for *S* = 0 mm. **c.** Effect of *d*_*o*_, *L*_*n*_ and *L*_*a*_ on *η* for 2 cm porcine tissue support with *S* = 0 mm.

For 100 *µ*l water injection in human skin supported by glass slab, a large variation in percentage delivery was observed (*η* ≈ 20 →95%). Whereas injections in human skin supported by porcine tissue yielded higher efficiency (*η >* 60%). Since applied loading is known to alter the structure and distribution of the forces within the skin layers, we used three different types of loading mechanisms (*A, B* and *C*). Type *B* and type *C* loading mechanism leads to high injection efficiency (*η >*80%) with small intrasample variation. However, skin stays under compression for longer time duration for loading mechanism type *B* and type *C*, which could challenge tissue integrity and, from the perspective of clinical implementation, are impractical. Standoff distance (*S* = 0 mm, 2 mm and 14 mm) was a highly significant factor for human skin injections, with *S* = 2 mm leading to complete delivery within the skin, followed by delivery efficiency (*η* ≈ 65-100%) obtained with water injected at *S* = 0 mm. Whereas, higher standoff distance of 14 mm showed lower injection efficiency (*η* ≈ 50-80%). The effect of ampoule volume was also significant, with nearly complete delivery of *V* = 50 *µ*l, but reduced efficiency for *V* = 100 *µ*l with large intra-sample variation. This indicates that there can be a limit in terms of deliverable volume (100*µl > V*_*c*_ ≥ 50*µ*l) for jet injection, which should not be exceeded in order to reduce wasted product and cross-contamination from rejected fluid.

Orifice diameter, viscosity of the injectate and the skin support showed significant effect on the delivery efficiency as presented in figure 10.B. The effect of injectate viscosity was very significant (*p <* 0.05) regardless of the skin support with an exception of injection via nozzle with *d*_*o*_ = 155 *µ*m into human skin supported by pork tissue (*p >* 0.05). Dyed water showed higher delivery into human skin in comparison to 80% glycerol injection. Also, effect of orifice exit diameter of nozzle on percentage delivery was not significant (*p >* 0.05) for different injectates and different skin supports.

The dual effect of applied load and orifice diameter is presented in figure 10.C, which indicates that a load of *L*_*n*_ ≈ 0.5 kg with an orifice diameter of 155 *µ*m provides the optimal configuration for loading mechanism type *A* (the most practical of the three) with *S* = 2 mm. As with GP skin, addition of axial skin tension is deleterious when used in combination with applied load.

## CONCLUSIONS

In this study we examined jet injection into freshly excised skin and skin that had undergone freeze-thaw cycle. One of the key factors for this study was that of applied load, as well as device parameters such as orifice diameter, standoff distance, ejected volume and fluid viscosity.

The bleb (fluid distribution pattern) formed in the intradermal tissues showed some qualitative variation in terms of overall shape, but across a wide range of parameters, we did not find statistically significant effects. As such, the principal metric we focused on was delivery efficiency, i.e. the ratio of liquid deposited under the skin to that ejected from the injector.

Nearly complete delivery of water was obtained for GP skin supported by lean pork when a normal load of 1 kg was applied. We hypothesize that this load represents a near-optimal load that removes the compliance of the underlying tissue, and correctly tensions the skin, but does not result in excessive compressive stress within the intradermal tissues. This is supported by results at higher normal loads (*L*_*n*_ = 2 kg) or those with added axial load, which showed reduced efficiency.

Anatomical variation of mechanical properties of skin was best highlighted using porcine skin, where we performed injections into cadavers, as well as freshly excised skin. In both cases, the highest injection efficiency occurred into ventral region skin, and we noted that injections with *V* = 50 *µ*l in fresh skin also achieved near-complete delivery (≈ 100%). After a single freeze-thaw cycle, same configuration performed poorly with lower percentage delivery of *η <* 55%. Other configurations exhibited nearly similar or slightly higher efficiency after a single freeze-thaw cycle, which we attribute to local structural damage caused by ice crystals.

In human skins, we used different loading protocols, and found that a two-cycle loading (type *B*) showed higher delivery efficiency. Increasing standoff to 14 mm showed low delivery efficiency. As with porcine skins, a lower injectate volume (*V* = 50 *µ*l) showed nearly complete delivery into the human skin as compared to 100 *µ*l, where the percentage delivery varied from ≈60% to ≈100%. Additional axial loading also led to poor injection efficiencies for human skins.

In summary, applied load, liquid properties, nozzle geometry and mechanical properties of the skin can strongly affect the delivery via jet injections, and the principal effect of these parameters are shown in table 1. Furthermore, optimal configurations for different skins are presented in table 2, which shows that *S* = 0 mm, *d*_*o*_ = *O*(100) *µ*m, *L*_*n*_ = 1 kg, *V* = 50 *µ*l is an all-round optimal configuration, other factors notwithstanding. These findings can be used to help guide future development of jet injectors.

**Table 1:** Summary of effect of parameters on delivery efficiency of the jet injections (*Decreases after *L*_*n*_ = 1 kg).

|  | Guinea Pig skin | Porcine skin | Human skin |
| --- | --- | --- | --- |
| **S, mm** | ↓ | ↓ | ↑↓ |
| $d_o$, $\mu\text{m}$ | — | — | ↑ |
| $L_n$, kg | ↑* | — | ↑ |
| $L_a$, kg | ↓ | — | ↓ |
| $V$, $\mu\text{l}$ | — | ↓ | ↓ |

**Table 2:** Optimal configuration for jet injections in different skin models.

|  | Guinea Pig skin | Porcine skin | Human skin |
| --- | --- | --- | --- |
| **S, mm** | 0 | 0 | 2 |
| $d_o$, $\mu\text{m}$ | 155 | 155 | 155 |
| **Loading mechanism** | — | — | $B$ |
| $L_n$, kg | 1 | 1 | 1 |
| $L_a$, kg | 0 | — | 0 |
| $V$, $\mu\text{l}$ | 50 | 50 | 50 |
| **Skin support** | 2 cm thick lean pork | — | 2 cm thick lean pork |

## ACKNOWLEDGMENTS

We would like to thank Department of the Animal Sciences at Texas Tech for porcine cadavers. This work was financially supported by Inovio Pharmaceuticals and The National Science Foundation via award CBET-1749382.

## AUTHOR CONTRIBUTIONS

**P.R.** and **J.M.** designed the experiments. **A.L.B., I.L.** and **P.R.** performed the experiments on the guinea pig skin. **VOB, C.W., W.T., I.L.** and **P.R.** performed the experiments on the human skin. **C.W., E.K. I.L.** and **J.M.** did the experiments on the porcine cadavers and excised the skin from the cadavers. **YR** contributed in experiments on the GP skin with 2 kg normal static load. **P.R.** analyzed all the data and wrote the manuscript with **J.M.**

