## Supplementary Info for "Loading effects on the performance of needle free jet injections in different skin models"

(Date: December 30, 2019)

### I. SUPPLEMENTARY VIDEOS

**Video 1:** Water injection in guinea pig skin supported by 2 cm thick pork meat, with a standoff distance of 14 mm and an applied normal load of 1 kg. *Video slowed down by 1000×.*

**Video 2:** Water injection in human skin supported by 2 cm thick pork meat, with a standoff distance of 14 mm and an applied normal load of 1 kg. *Video slowed down by 1000×.*

### II. HUMAN SKINS CATALOG

All the human skins were procured from the National Disease Research Interchange (NDRI, PA) and were non-reactive (*negative for following infectious diseases: HIV I & II, Hep B antigen and Hep C antibody*). Skins were kept in a freezer at a temperature of -20°C and were thawed to room temperature (20°C) before performing jet injections.

Table S1: Human skins used in the study

| S. No. | Age | Race | Sex | H/W (in/lbs) | Location |
|---|---|---|---|---|---|
| 1 | 67 | Caucasian | Female | 62/218 | Left Deltoid |
| 2 | 75 | Caucasian | Female | -/- | Left Deltoid |
| 3 | 80 | Caucasian | Male | 68/161 | Left Deltoid |
| 4 | 55 | Caucasian | Male | 74/281 | Left Deltoid |
| 5 | 72 | Caucasian | Male | 72/220 | Right Deltoid |
| 6 | 74 | Caucasian | Male | 73/163 | Left Deltoid |
| 7 | 66 | Caucasian | Female | 62/152 | Abdomen |
| 8 | - | - | - | - | - |
| 9 | - | - | - | - | - |

### III. CHARACTERIZATION OF LIQUID JET

Nozzle orifice with orifice exit diameter ($d_o$) of 155 $\mu$m and 175 $\mu$m were used. Liquid jet speed at the nozzle exit was estimated by tracking plunger displacement. Plunger speed was then converted to jet exit speed ($v_j$) using equation of mass conservation.

Jet speed for dyed water and dyed 80% glycerol for nozzle with nozzle orifice diameter of 155 $\mu$m was 143.45±2.69 m/s and 121.69±1.57 m/s respectively. Increasing $d_o$ to 175 $\mu$m gives jet speed of 124.01±2.66 m/s and 102.53±2.38 m/s for dyed water and dyed 80% glycerol respectively.

Table S2: Liquid injectate properties

| **Liquid** | $\mu$ (mPa.s) | $\rho$ (kg/$m^3$) |
|---|---|---|
| Water | 1 | 994.84 |
| 80% Glycerol | 84 | 1201.61 |
| Water + Dye | $\sim$ 1 | 996.07 |
| 80% Glycerol + Dye | $\sim$ 84 | 1202.32 |

### IV. INJECTION DELIVERY EFFICIENCY

Delivery efficiency of water and 80% glycerol with variation in normal load ($L_n$) showed similar trend for guinea pig skin supported by 1 cm thick lean pork. On contrary, jet injection delivery in guinea pig skin supported by pork fat layer showed no significant effect of applied normal load for water and 80% glycerol without any standoff distance.

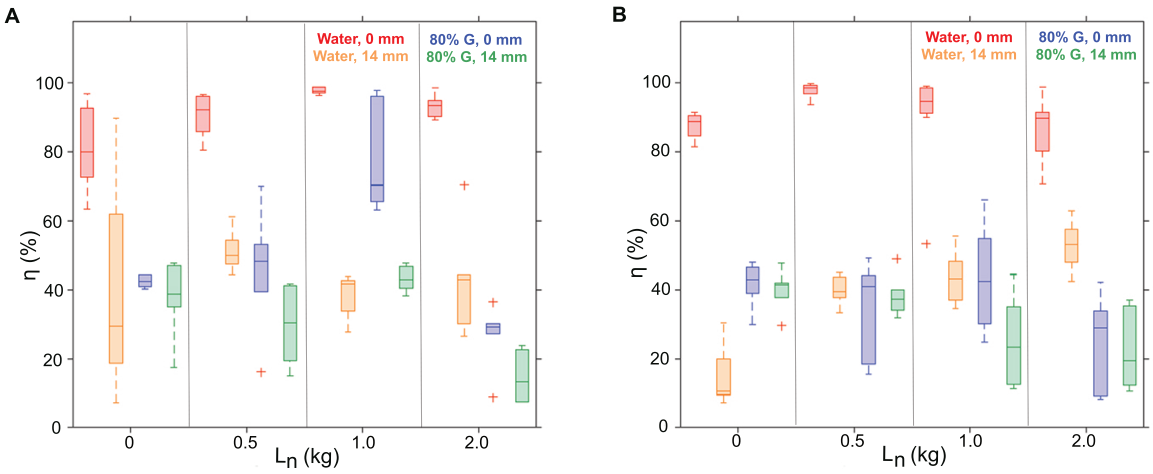


Figure S1: **Injection efficiency ($\eta$) of jet injections in guinea pig skin. Effect of normal load ($L_n$), standoff distance ($s$) and injectate viscosity ($\mu$) on $\eta$ for: a.** GP skin supported by 1 cm thick leaned pork **b.** GP skin supported by 1 cm thick fat layer

### V. BLEB DIMENSIONS

Bleb dimensions were measured via image processing in Matlab 2017b. Aspect ratio (AR) was used as a parameter to understand the effect of various parameters on the dispersion pattern formed by the liquid inside the skin.

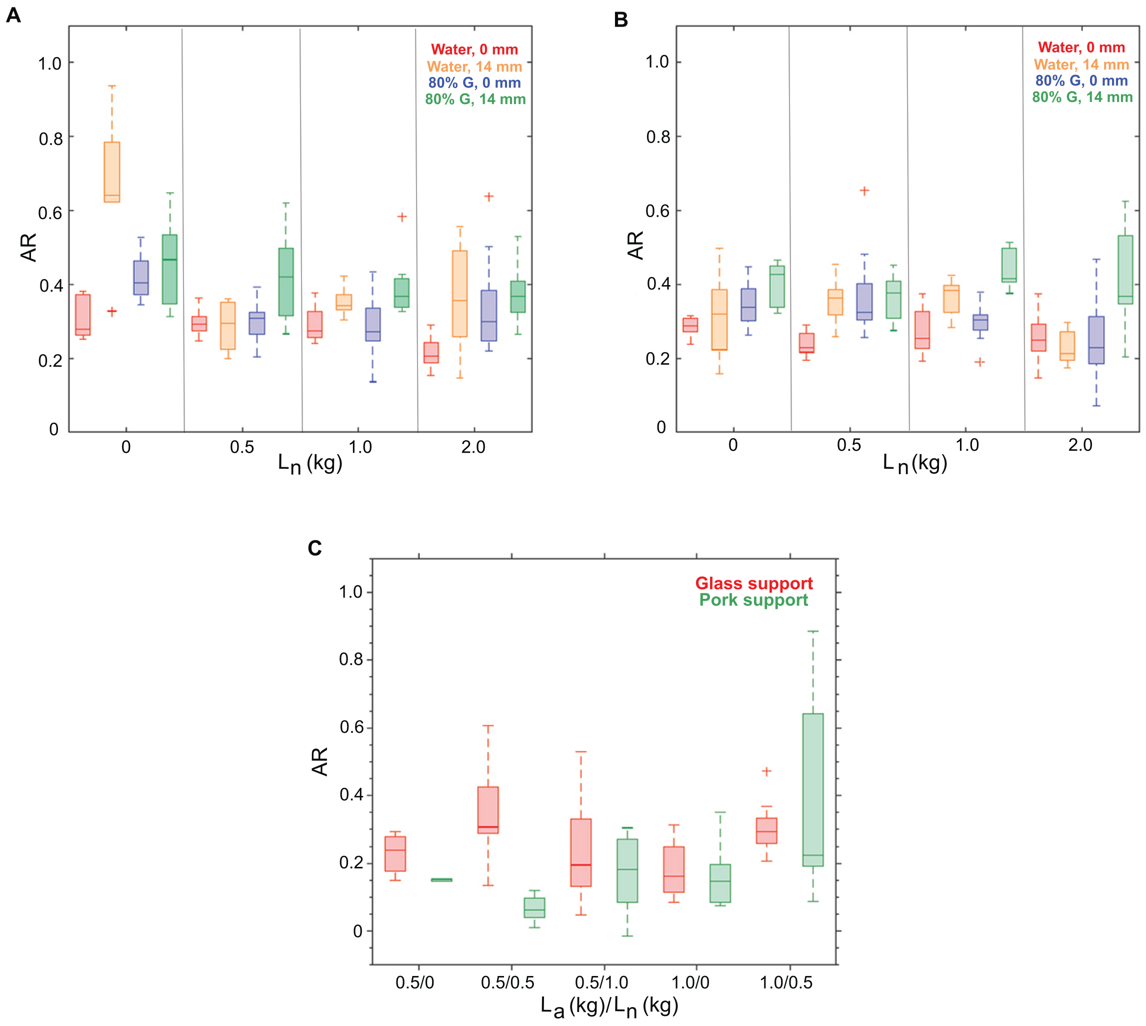


Figure S2: **Aspect ratio (AR) of blebs in guinea pig skin. Effect of normal load ($L_n$), standoff distance ($s$) and injectate viscosity ($\mu$) on AR for: a.** Skin supported by 1 cm thick leaned pork **b.** Skin supported by 1 cm thick fat layer. **c. Effect of axial loading on bleb dimensions in guinea pig skin**

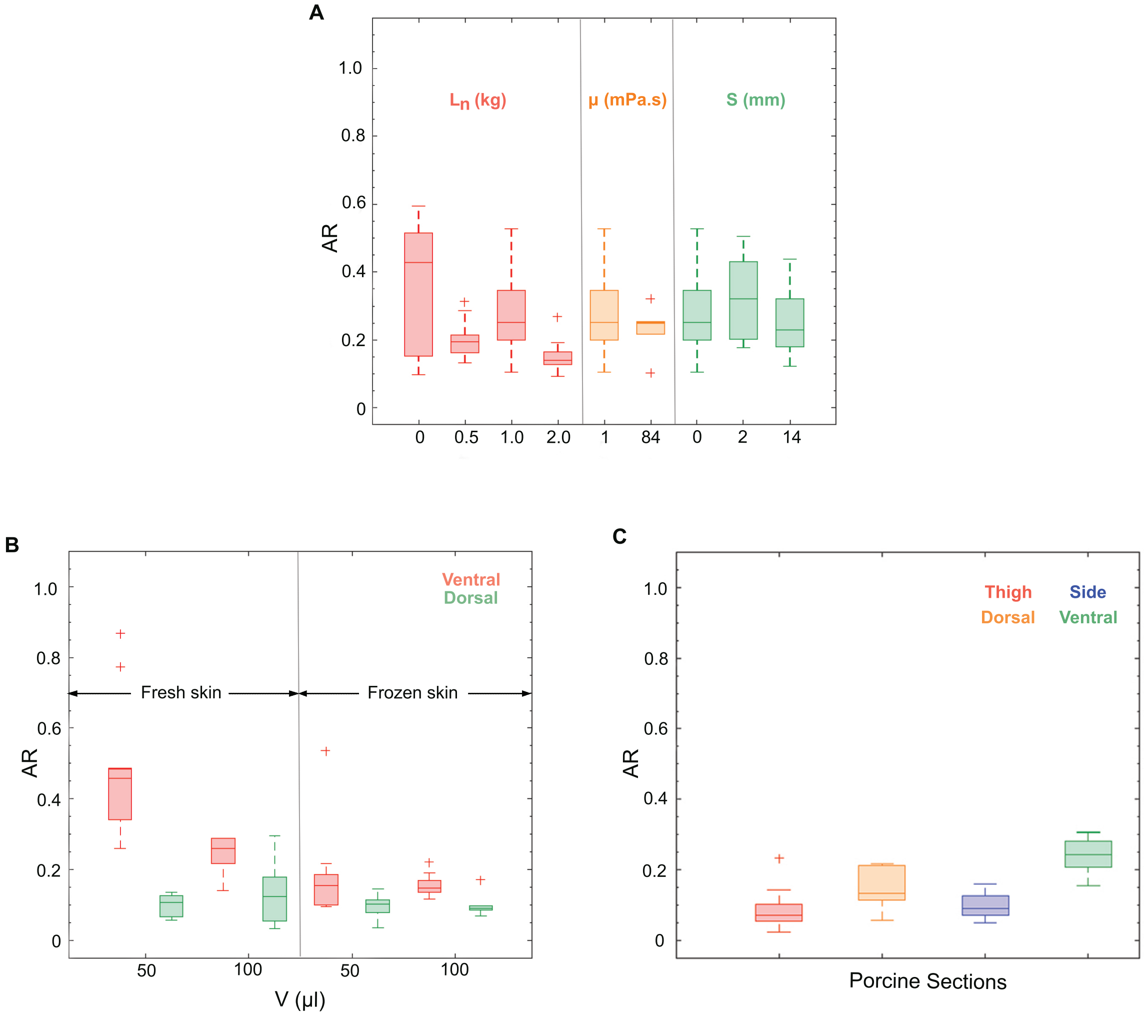


Figure S3: **Aspect ratio of the blebs obtained for the jet injections in porcine skin a.** Effect of applied normal load ($L_n$, $kg$), injectate viscosity ($\mu$, $mPa.s$) and standoff ($s$, $mm$) on the AR of the blebs formed in the fresh excised porcine skin (* represents $D_o$ = 175 $\mu$m, otherwise $D_o$ = 155 $\mu$m). **b.** AR of the blebs in the porcine skin from ventral and dorsal regions affected by the state of the skin, thickness and liquid volume after a single freeze and thaw cycle. **c.** Aspect ratio of the blebs formed in the skin in the porcine cadaver into thigh, ventral, side and dorsal regions.

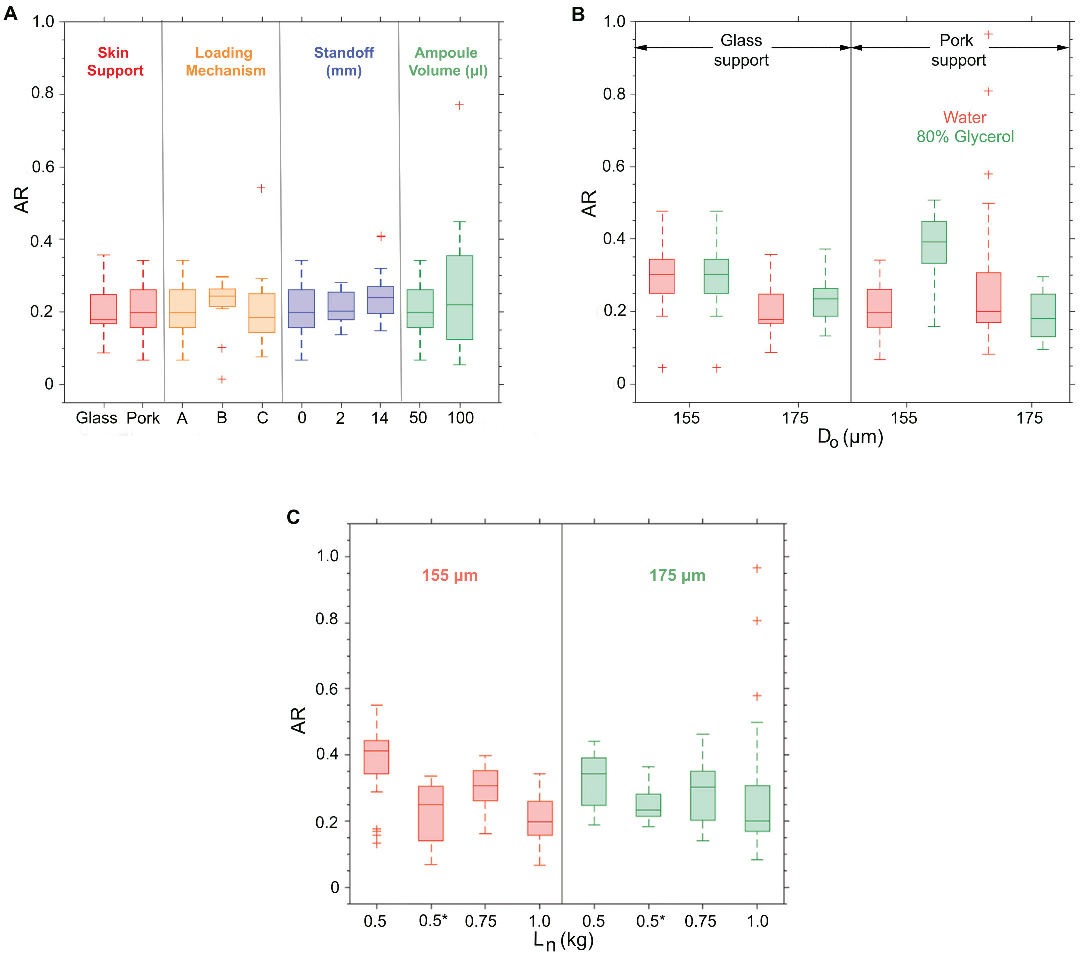


Figure S4: **Aspect ratio of the blebs after jet injections in human skin**. **a.** Effect of skin support, loading mechanism, standoff and V on AR of the blebs formed in the skin for $L_n = 1$ kg. **b.** Effect of $D_o$, $\mu$ and skin support on bleb AR for $L_n = 1$ kg. **c.** Effect of $D_o$, $L_n$ and $L_a$ on bleb AR. (**represents additional axial load of 0.5 kg using hooked weights*)

### VI. EVAPORATION FROM SKINS DURING EXPERIMENTS

The water content of different skins was observed to diminish slowly due to evaporation. Thus, estimation of delivery efficiency was unreliable on the basis of weight of the skin before and after the jet injection. Therefore, percentage delivery of different liquids was measured on the basis of the rejected liquid absorbed by the filter paper. Skin was weighed before and after the injection, with three measurements for each case in addition to recording the time duration of the measurements. Time and weight recordings were used to measure the evaporation rate of water content from different skins in mg/min. The water content of skin is an important parameter which can alter mechanical properties of the skin. It is worth noting that skins were thawed to room temperature before every experiment without any humidity control.

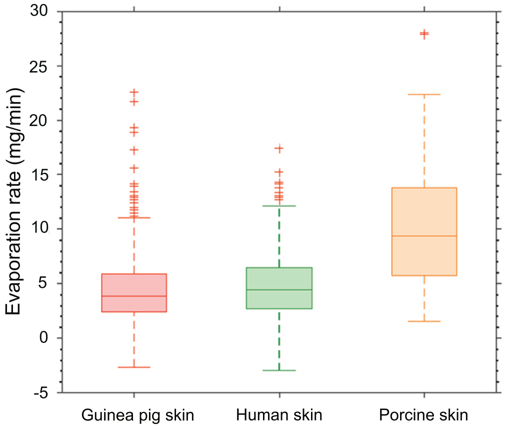


Figure S5: **Evaporation rate from different skins**

Guinea pig and human skins were procured in frozen state, while porcine skin was obtained from cadavers. Thus, water content of porcine skin (*not quantified*) was different from that of the guinea pig skin and human skin. Different water content possibly leads to different evaporation rates for different skins as observed in figure S5.

### VII. SKIN UNDER TENSION BY SPACER RING

Figure S6 shows how a spacer ring attached to the nozzle of a jet injector deforms the skin and creates tension on the skin for an applied normal load of 1 kg. The spacer ring serves dual purpose of generating tension on the skin and providing a standoff distance between the nozzle exit orifice and the skin.

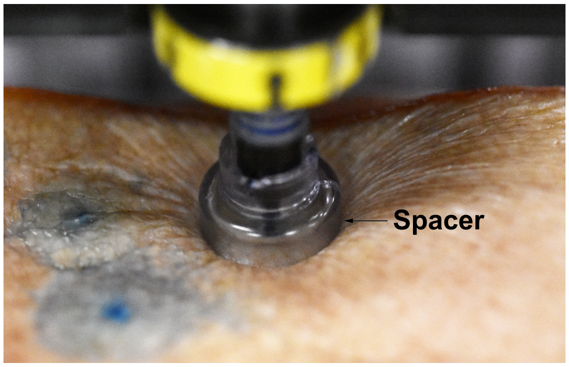


Figure S6: **Human skin under tension by a spacer ring with an applied load of 1 kg by jet injector.**

### VIII. ANOVA ANALYSIS

#### A. GUINEA PIG SKIN

Table S3: p-value: Effect of applied load in guinea pig skin

| | **Leaned pork, 1 cm** | **Leaned pork, 2 cm thick** | **Fat, 1 cm** |
|---|---|---|---|
| **Water, s = 0 mm** | 0.025 | 0.0012 | 0.3638 |
| **Water, s = 14 mm** | 0.718 | 0.2482 | $3.4\times10^{-6}$ |
| **80% Glycerol, s = 0 mm** | 0.0002 | 0.0055 | 0.2514 |
| **80% Glycerol, s = 14 mm** | 0.0027 | 0.177 | 0.0125 |

Table S4: p-value: Effect of skin support in guinea pig skin

| | $L_n$ = **0 kg** | $L_n$ = **0.5 kg** | $L_n$ = **1 kg** | $L_n$ = **2 kg** |
|---|---|---|---|---|
| **Water, s = 0 mm** | 0.5926 | 0.145 | 0.2938 | 0.0455 |
| **Water, s = 14 mm** | 0.1613 | 0.0013 | 0.009 | 0.0288 |
| **80% Glycerol, s = 0 mm** | 0.9157 | 0.4858 | 0.0124 | 0.2736 |
| **80% Glycerol, s = 14 mm** | 0.4312 | 0.3566 | 0.069 | 0.0072 |

Table S5: p-value: Effect of standoff distance in guinea pig skin

| | $L_n$ = **0 kg** | $L_n$ = **0.5 kg** | $L_n$ = **1 kg** | $L_n$ = **2 kg** |
|---|---|---|---|---|
| **Water, Leaned pork (2 cm)** | 0.0262 | $5.347\times10^{-6}$ | $7.06\times10^{-11}$ | $2.97\times10^{-5}$ |
| **Water, Leaned pork (1 cm)** | 0.0295 | $1.07\times10^{-5}$ | $4.46\times10^{-8}$ | 0.002 |
| **Water, Fat (1 cm)** | 0.0117 | $2.79\times10^{-7}$ | 0.0007 | 0.0015 |
| **80% G, Leaned pork (2 cm)** | 0.9236 | 0.2759 | 0.0201 | 0.0001 |
| **80% G, Leaned pork (1 cm)** | 0.2294 | 0.2085 | 0.0017 | 0.1014 |
| **80% G, Fat (1 cm)** | 0.4229 | 0.8182 | 0.821 | 0.2956 |

Table S6: p-value: Effect of viscosity in guinea pig skin

| | $L_n$ = **0 kg** | $L_n$ = **0.5 kg** | $L_n$ = **1 kg** | $L_n$ = **2 kg** |
|---|---|---|---|---|
| **s = 0 mm, Leaned pork (2 cm)** | 0.0012 | 0.019 | 0.0038 | $5.26\times10^{-6}$ |
| **s = 14 mm, Leaned pork (2 cm)** | 0.7598 | 0.5598 | 0.0288 | 0.0511 |
| **s = 0 mm, Leaned pork (1 cm)** | 0.0002 | 0.0013 | 0.0341 | $9.155\times10^{-7}$ |
| **s = 14 mm, Leaned pork (1 cm)** | 0.7745 | 0.0127 | 0.1873 | 0.0131 |
| **s = 0 mm, Fat (1 cm)** | $1.263\times10^{-6}$ | $1.344\times10^{-5}$ | 0.0046 | $7.92\times10^{-5}$ |
| **s = 14 mm, Fat (1 cm)** | 0.0017 | 0.7323 | 0.059 | 0.0009 |

Table S7: p-value: Effect of normal static load in addition to axial load in guinea pig skin

| **Glass, $L_a$ = 0.5 kg** | **Glass, $L_a$ = 1 kg** | **Pork, $L_a$ = 0.5 kg** | **Pork, $L_a$ = 1 kg** |
|---|---|---|---|
| 0.6058 | 0.0104 | 0.0028 | 0.0195 |

Table S8: p-value: Effect of skin support for applied normal static load in addition to the axial loading in guinea pig skin

| $L_a$ = **0.5 kg,** $L_n$ = **0 kg** | $L_a$ = **0.5 kg,** $L_n$ = **0.5 kg** | $L_a$ = **0.5 kg,** $L_n$ = **1 kg** | $L_a$ = **1 kg,** $L_n$ = **0 kg** | $L_a$ = **1 kg,** $L_n$ = **0.5 kg** |
|---|---|---|---|---|
| 0.3217 | $8.83\times10^{-7}$ | $5.38\times10^{-5}$ | 0.9927 | 0.0408 |

### B. PORCINE SKIN

Table S9: p-value: Effect of various parameters in porcine skin for dyed water injection

| **Anatomy** | $L_n$ | **Standoff** |
|---|---|---|
| 0.004 | 0.946 | 0.135 |

Table S10: p-value: Effect of injectate viscosity

| $L_n$ = **1 kg** |
|---|
| 0.068 |

### C. HUMAN SKIN

Table S11: p-value: Effect of different parameters for dyed water injection in human skin with $L_n = 1$ kg

| **Skin support** | **Loading mechanism** | **standoff distance** | **ampoule volume** |
|---|---|---|---|
| 0.203 | 0.875 | 0.171 | 0.811 |

Table S12: p-value: Effect of liquid viscosity in human skin with $L_n = 1$ kg for pork support

| $d_o = \mathbf{155}\ \mu\mathbf{m}$ | $d_o = \mathbf{175}\ \mu\mathbf{m}$ |
|---|---|
| 0.3909 | 0.0005 |

Table S13: p-value: Effect of liquid viscosity in human skin with $L_n = 1$ kg for glass support

| $d_o = \mathbf{155}\ \mu\mathbf{m}$ | $d_o = \mathbf{175}\ \mu\mathbf{m}$ |
|---|---|
| 0.0036 | 0.0075 |

Table S14: p-value: Effect of $D_o$ in human skin on glass support with $L_n = 1$ kg

| $\mu = \mathbf{1\ mPa.s}$ | $\mu = \mathbf{84\ mPa.s}$ |
|---|---|
| 0.827 | 0.455 |

Table S15: p-value: Effect of $D_o$ in human skin on pork support with $L_n = 1$ kg

| $\mu = \mathbf{1\ mPa.s}$ | $\mu = \mathbf{84\ mPa.s}$ |
|---|---|
| 0.615 | 0.080 |

Table S16: p-value: Effect of $L_n$ in human skin on leaned pork (2 cm) support for dyed water injection

| $D_o = \mathbf{155}\ \mu\mathbf{m}$ | $D_o = \mathbf{175}\ \mu\mathbf{m}$ |
|---|---|
| 0.99 | 0.39 |

Table S17: p-value: Effect of $L_a$ in human skin on leaned pork (2 cm) support for dyed water injection with $L_n = 0.5$ kg

| $L_a = \mathbf{0.5\ kg}$ |
|---|
| 0.097 |
